# Genome Scale-Differential Flux Analysis reveals deregulation of lung cell metabolism on SARS Cov2 infection

**DOI:** 10.1101/2020.11.29.402404

**Authors:** Piyush Nanda, Amit Ghosh

## Abstract

The COVID-19 pandemic is posing an unprecedented threat to the whole world. In this regard, it is absolutely imperative to understand the mechanism of metabolic reprogramming of host human cells by SARS Cov2. A better understanding of the metabolic alterations would aid in design of better therapeutics to deal with COVID-19 pandemic. We developed an integrated genome-scale metabolic model of normal human bronchial epithelial cells (NHBE) infected with SARS Cov2 using gene-expression and macromolecular make-up of the virus. The reconstructed model predicts growth rates of the virus in high agreement with the experimental measured values. Furthermore, we report a method for conducting genome-scale differential flux analysis (GS-DFA) in context-specific metabolic models. We apply the method to the context-specific model and identify severely affected metabolic modules predominantly comprising of lipid metabolism. We conduct an integrated analysis of the flux-altered reactions, host-virus protein-protein interaction network and phospho-proteomics data to understand the mechanism of flux alteration in host cells. We show that several enzymes driving the altered reactions inferred by our method to be directly interacting with viral proteins and also undergoing differential phosphorylation under diseased state. In case of SARS Cov2 infection, lipid metabolism particularly fatty acid oxidation and beta-oxidation cycle along with arachidonic acid metabolism are predicted to be most affected which confirms with clinical metabolomics studies. GS-DFA can be applied to existing repertoire of high-throughput proteomic or transcriptomic data in diseased condition to understand metabolic deregulation at the level of flux.

## Introduction

In several disease conditions, metabolic reprogramming serves as the epicenter for loss of homeostasis in the host cell(1–4). Metabolism allocates biochemical resources to the cellular machinery. Disease progression either caused by internal disturbances in host cell or attack by pathogenic intruders require reallocation of biochemical resources(5, 6). Apart from resource synthesis, metabolism has also been shown to reciprocally interact with other molecular events like epigenetic modifications and gene regulation(7–10). The role of metabolism in several high incidence diseases like cancer(11, 12), tuberculosis(13) and neuronal disorders(14) have been elucidated in fine details with the advent in high-throughput technologies to study metabolites and underlying reactions. Therefore, technological and methodological development in understanding metabolic states of cells will help us in gaining better understanding of disease mechanisms and strategies to design better therapeutics.

In the context of the current COVID-19 pandemic, several reports highlight the deregulation of multiple metabolic pathways in the host cell in response to SARS Cov2 infection(15–17). Cases of altered lipid metabolism is observed in patients who were infected with SARS Cov between 2002 and 2003, 12 years after their recovery. Moreover, reports of altered tryptophan and lipid metabolism has been elucidated in clinical samples of COVID-19 patients. Understanding the metabolic alterations at a genome-scale will help us in gaining deeper insights into the mechanism of pathogenesis beyond immunological effects. With increasing evidence on the importance of metabolism in disease biology in particular the current COVID-19, it has become imperative to develop technologies to probe into activities of reactions that underlie metabolic pathways. Flux, which is defined as the reaction rate per unit biomass, directly reflects on the homeostatic balance of the cell. Disturbance of cellular homeostasis due to pathogenic intruders is clearly reflected in alterations in metabolic pathways. In this regard, Genome-Scale Metabolic Models (GEMs) have been leveraged to understand flux distributions(18–20) using tools like Flux Balance Analysis (FBA). Through years, GEMs of human cell lines have been used for understanding progression of disease, host-pathogen interactions and up to neurological disorders. Algorithms like GIMME, iMAT, MBA, INIT/tINIT have enabled us to incorporate gene expression or proteomics data into the model (GEMs) and hence furnishing tissue or cell or condition specific GEMs(21). This enables us to enlist reactions which are differentially altered between normal and diseased state. Highly curated Human GEMs like Recon3D(22) and HumanGEM(23) have incorporated finer details about the human metabolism. They have been validated under multiple experimental conditions and have shown to be representative of the human metabolism in both normal and diseased conditions.

Despite the advancement in metabolic modeling, most of the work has focused on elucidating altered reactions based on differentially incorporated reactions in condition-specific metabolic models. Such an approach, allows us to map the effects on gene expression or enzyme production on reactions qualitatively. Our ability to find the degree of alterations at the level of fluxes in such context specific models will enable us to understand the activities of different pathways in the cell. Fluxes are a consequence of interactions between several reactions, the structural features of the metabolic pathways and the nutrient uptake rates(24). Limited analysis has been done to leverage the power of metabolic flux analysis to probe into differences in fluxes between diseased and non-diseased state. The altered reactions calculated using the differences in flux would also enable us to conduct enrichment analysis of pathways and pin point specific metabolic modules altered between two conditions i.e. normal and diseased. Although computational methods exists to infer metabolic changes using relative gene expression(25) or transcriptome data(26) data but none infer differences at the level of flux directly. FBA, while being a powerful tool in microbial systems biology, requires a strict objective function such as specific growth rate(27). This restricts its usage in the case of GEMs of higher eukaryotes where it’s difficult to define single cellular objectives. Approaches like Flux Variability Analysis (FVA)(28) and flux sampling(29) alleviate this problem by allowing us to obtain the limits of the solution space. These tools have not been put into a statistical framework to allow us to conduct differential flux analysis and pathway enrichment studies as we do for gene expression.

While such steady state approaches, allows us to predict reaction fluxes under given conditions, it doesn’t speak much about the mechanism of flux regulation. To understand the regulation of reactions, it becomes imperative to conduct integrative analysis of metabolic flux analysis with other omics approaches. Omics datasets such as protein-protein interaction networks and phosphorylation variations can help us in inferring the prospective causative factors behind alterations in metabolic flux. Therefore, our ability to develop pipelines for integrated analysis of drivers (alterations in post-translational modifications of enzymes) and effects (fluxes) will enable us to obtain better insights into disease mechanisms.

In this study, we developed a Genome-Scale Differential Flux Analysis (GS-DFA) that leverages flux sampling and stringent statistical framework to report affected metabolic modules in diseased conditions. We developed a context-specific integrated GEM of SARS Cov2 infected NHBE cells leveraging the publicly available gene expression data. We apply the GS-DFA to the context specific GEM of normal human lung cells or tissues (NHBE and Lung Biopsy) and SARS Cov2 infected human lung cells. We show that several expected pathways are affected and significant number of altered pathways have associations with viral proteins through PPIs or/and altered phosphorylation landscape. Several pathways such as fatty acid oxidation, fatty acid biosynthesis and amino acid metabolism is affected in diseased state. Such pathways are also associated with the production of immune-regulatory molecules like Leukotriene and Prostaglandins which mediate cell-to-cell signaling and inflammatory response management(30). SARS Cov2 proteins interacts with multiple enzyme subunits driving the reactions we enlisted to be differentially altered. This indicates a mechanism by which the virus possibly reprograms the metabolism of the host cell in order to facilitate its growth and survival.

## Results

### Stoichiometric model of SARS Cov2 biomass

Flux balance analysis requires a biomass equation which would serve as an objective function for the underlying linear optimization problem(24). The biomass equation describes the accurate stoichiometry of the biological system in question based on the molecular make up of its constituent proteins, nucleic acids, carbohydrates and lipids. We sought to define a similar biomass equation for SARS Cov2. Integrating the same into the genome-scale metabolic model of the human lung cells would allow us to analyze the host-virus metabolic interaction. Each virus particle comprises of a single stranded positive sense RNA, proteins at the membrane and in complex with the RNA, carbohydrates studded on some of the membrane proteins and a lipid bilayer(31) (Figure 1A). Leveraging the recent literature data on copy number of the macromolecules and their chemical composition, we constructed a virus biomass objective function (or biomass equation)-Cov2VBOF. We used spike-protein copy number per virion as a basis to calculate the absolute copy number of proteins on the virus leveraging proteomics datasets. Here, the fractional molecular contribution of each amino acid was calculated solely from the copy number of proteins and their amino acid composition (Figure 1B). Similarly, we used information on translational efficiency over the viral genome(32) and the absolute copy number of the proteins, to calculate the mRNA levels resulting from the viral genome. Several studies show the low GC content of the virus is intrinsically linked to a strategy to avoid recognition by the innate immune system. Particularly, ZAP proteins recruit themselves to CpG enriched regions and trigger host machinery to degrade the viral genome(33). We therefore observe a lower fraction of GC in the SARS Cov2 genome and the corresponding transcriptome (Figure 1C).

**Figure 1.**
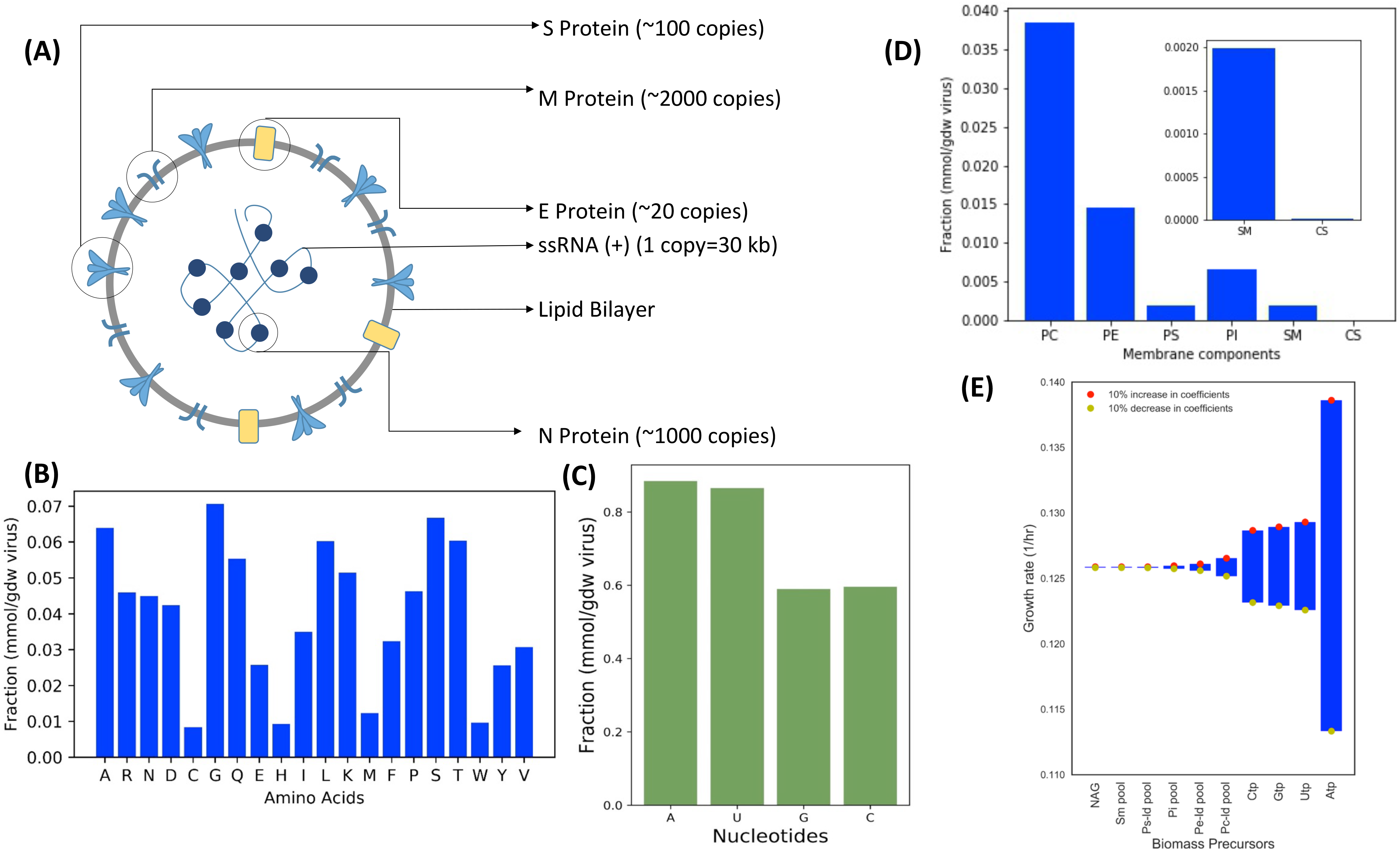
Reconstruction of biomass equation (VBOF) of SARS Cov2 virus from its stoichiometric make-up. (A) The stoichiometry of various proteins, nucleic acid and their locations in the SARS Cov2 virus. The molar composition of constituent molecules in lipid fraction (B), nucleic acid (C) and proteins (D). PC-Phosphatidylcholine; PE-Phosphatidylethanolamine; PS-Phosphatidylserine; PI-Phosphatidylinositol; SM-Sphingomyelin; CS-Cholesterol. (E) Sensitivity analysis of growth rate of SARS Cov2 with respect to coefficient (fractional composition) of biomass precursor. The coefficients are varied by ±10% and y-axis shows growth rates simulated by Flux Balance Analysis (FBA).

Phosphatidylcholine and Phosphatidylethanolamine occupied the highest lipid molar fraction in the membrane whereas cholesterol occupied the lowest molar fraction (Figure 1D). The biomass equation has 44 biomass components including all 20 amino acids, 4 nucleotides, lipid and cholesterol, carbohydrates along with ATP required for polymerization of the macromolecules. N-Acetyl Glucosamine (NAG) is a molecular coating on the Spike protein of SARS Cov2 which allows the virus to mask the potential epitopes against which the host cell can develop anti-bodies(34). The SARS Cov2 infection causes activates the NAG biosynthesis pathway in order to meet the viral demands. To show the importance of biomass precursors towards fitness of the virus, we conducted a biomass sensitivity analysis. Here, we tried to capture the effect of small fluctuations (~10%) in coefficient of biomass precursor on the specific growth rate prediction by FBA (Figure 1E and Figure S4). We show that the nucleotides (highest), lipids and NAG (lowest but non-zero) affect the specific growth rate the most. Such an analysis allows us to rank the biomass precursors based on their relative importance on the fitness. As a significant fraction (Figure 1D) of SARS Cov2 biomass is based on its nucleic acid, it is likely that fluctuations in their components will affect the growth rate the most. The components of lipid membrane also affect the growth rate of the virus significantly as unlike amino acids, they are de novo synthesized in the host cell. Similarly, N-acetylglucosamine is synthesized de novo to serve as a carbohydrate coating on the surface proteins of SARS Cov2. This analysis allows us to understand the biomass components of SARS Cov2 whose variation might affect the growth rate of SARS Cov2 severely.

We further integrated the Cov2VBOF into the context specific models of NHBE cells in infected state. We showed the Cov2VBOF helps in accurately simulating the growth kinetics of the virus upon infection. By varying the coefficient of each precursor by 50% (an extreme variation), we were able to show the predicted growth rate shows no significant difference with the experimental reported growth rates of the virus (p>0.05, t-test) (Figure 2C). As several different methods were used to measure the experimental growth rate(35) in literature, the constructed biomass equation for SARS Cov2 confirms with the realistic chemical composition of the virus.

**Figure 2.**
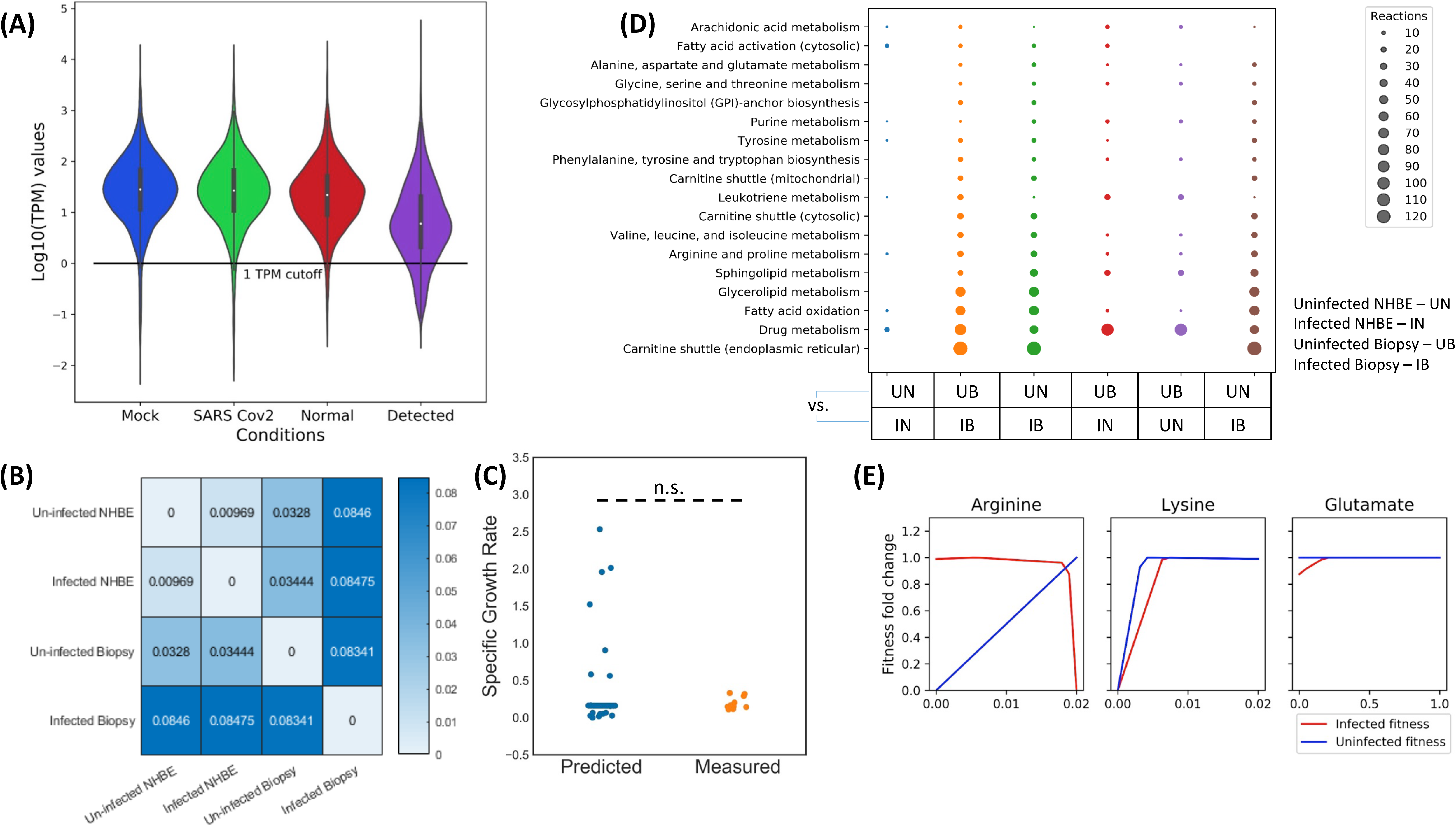
Generation of context-specific models of SARS Cov2 infected cells and normal cells using gene-expression data and tINIT algorithm. (A) The distribution of TPM normalized gene expression in Mock NHBE cells, SARS Cov2 infected NHBE cells, Normal Lung Tissue (From Biopsy) and Infected Tissue (From Biopsy). (B) The Hamming distance between all the generated context specific models based on the differences in reactions in the model. (C) The agreement of model simulated specific growth rate of the virus with the experimental reported specific growth rate. n.s. stands for non-significant differences as estimated by t-test. (D) The pathways over-represented while comparison between models. The reactions over-represented in each comparison are present exclusively in one of the models. (E) The robustness of specific growth rate of SARS Cov2 with respect to changes in specific uptake rates of arginine, lysine and glutamate. Refer to Figure S3 for robustness analysis with respect to specific uptake rates of all nutrients.

### Integration of transcriptomic profile in Human GEM

In the case of mammalian cells, it becomes particularly important to constrain the reactions in the generalized HumanGEM metabolic model based on expression levels in the cells to define the metabolic model of a given cell type. This eliminates reactions that are lowly expressed in the specific cell/tissue while keeping reactions having high expression or are required for basic metabolic functionality. We used task based INIT(36) (Integrative Network Inference for Tissues) to define NHBE cell and Lung Biopsy context specific metabolic models with and without viral infection. The TPM normalized gene expression data(37) (Figure 2A) was integrated into the model as described in material and methods. The NHBE background context specific metabolic models were named as iNHBE and iNHBECov2 for normal NHBE and infected NHBE respectively. Similarly, the lung biopsy specific metabolic models were named iLungsTissue and iLungsTissueCov2. Majority of altered reactions in infected NHBE cells (iNHBECov2) compared to normal NHBE (iNHBE) belonged to arachidonic acid metabolism, fatty acid oxidation, fatty acid activation and few amino acid biosynthesis pathways of arginine, proline and tyrosine (Figure 2D). Deregulation of arachidonic acid metabolism has been reported earlier to be associated with coronavirus infection. Linoleic acid to arachidonic acid metabolism axis is one of the most perturbed reactions as reported through metabolomics analysis of HCov-229E infected cells(38). We also observe a perturbation in Leukotriene pathways between the iNHBE and iNHBECov2 cells (Figure 2D). Arachidonic acid is the precursor for the production of leukotrienes and prostaglandins through oxidation(30). Leukotrienes are also important cell signaling molecules which are involved in both autocrine and paracrine signaling(30). More particularly, leukotrienes are important class of molecules that have chemotactic action of migrating immune cells. Prostaglandins are mediators of inflammatory responses in the cell(30). Similarly, between iLungsTissue and iLungsTissueCov2, apart from pathways affected as in the case of NHBE, we observed differences in reactions belonging to carnitine shuttle (endoplasmic reticulum and mitochondria), glycerolipid and sphingolipid metabolism (Figure 2D). There were also differences in several amino acid metabolism pathways of glycine, serine, alanine and aspartate. The differences in prediction of affected reactions in the case of NHBE cell context specific GEMs and Lung Biopsy context specific GEMs could be due to the heterogeneity of tissue samples. The affected reactions in the case of lung tissue biopsy presents the metabolic alterations at the level of tissue resulting from metabolic interactions among the constituent cell types. NHBE cells are laboratory adapted epithelial lung cells and the context specific GEMs corresponding to cellular level response to the infection. We further compared the context specific GEMs of NHBE cells and the lung tissue biopsy to elucidate fundamental differences in metabolic state both in infected and uninfected conditions. We used Hamming distance as a proxy for quantifying the differences in the metabolic architecture of the context specific models (Figure 2B). Carnitine shuttle pathway, which activates fatty acids and transports them for fatty acid beta-oxidation, is a major affected pathway in the case of lung tissue biopsy of infected patients compared to normal (Figure 2B). This pathway is not particularly observed to be an affected pathway in the case of NHBE cells. We hypothesize the differences might be due to metabolic interactions at the level of tissue and contribution of heterogeneous cell types.

The Cov2VBOF was leveraged to measure the specific growth rate of the virus in the context specific model of the infected NHBE cell. We used Flux Balance Analysis (FBA) on the context specific model constrained by HAM media using Cov2VBOF as an objective function. We repeated the FBA for several perturbations in the Cov2VBOF (to account for effect of uncertainty in biomass composition on growth rate) and small alterations in the uptake rates of nutrient in the infected conditions (Accounting for effect of uncertainties in uptake rates of media components). We compared the predicted specific growth rate to the experimental growth rate as reported in earlier(39). We could find no significant differences between the predicted and experimental growth rate (p>0.05, t-test) (Figure 2C). This is a strong evidence for the validity of the constructed virus biomass objective function.

Furthermore, we tried to capture the effect of specific uptakes rates (mmol/gdw/hr) of nutrients on the fitness of the virus in infected cells through phenotypic phase plane analysis. Such an analysis allows to visualize the effects of alterations in uptake rates (Phenotype 1) on the growth of the virus i.e. fitness (Phenotype 2) (Figure S1). It was interesting to note that increased uptake rates of arginine severely affects the fitness of the virus while it supplements the growth of the normal NHBE cells (Figure 2E). We also noted the effect of decreased uptake of lysine and glutamate on the reduction in fitness of the virus (Figure 2E). Amino acid transport in the cells are mostly driven by facilitated diffusion through transporters. Therefore, the concentration of amino acids in the cellular environment would dictate the uptake rate of amino acids. Similar approach of supplementing or depleting amino acids have been used as therapeutic option for treating of cancer and other diseases(40).

Context specific metabolic models of NHBE cells and lung biopsy helped us to catalogue altered metabolic reactions between normal and infected condition. The network properties of the metabolism and the gene-protein-reaction association captured by the HumanGEM model combined with the gene expression data allowed us to analyze altered metabolic phenotypes in diseased state. These models can be further leveraged to reveal more biological insights into the metabolic state of the diseased cell using powerful tools of metabolic flux or network analysis.

### Differential Flux Analysis for hunting altered reactions in SARS Cov2 infected cell

Integration of gene expression data into the highly curated HumanGEM pruned non-essential reactions while enriching important reactions. This helped us to generate context specific models of normal and diseased cells. We further aimed at understanding the alterations in metabolic flux in an infected cellular state. Such an analysis would pinpoint specific reactions which show altered behavior in diseased state. While inferring altered metabolic pathways have been conventionally done using differential gene expression analysis, we show the limitations of such analysis. The presence of either isozymes (multiple enzymes having same function) or multi-subunit complex (multiple subunits contributing collectively towards a function) makes it difficult to infer the effects of altered gene expression on metabolism(21). It is very well known that alterations in levels of enzyme facilitating the rate limiting step can lead to changes in flux distributions in the downstream reactions. In a hypothetical scenario (Figure 3A), we can see only the alterations in the expression of Gene A is sufficient to change the flux through Reaction 1 which further pushes the flux through Reaction 2. This is irrespective of changes in expression levels of Gene C and D. Therefore, to enlist affected reactions or pathways, it is imperative to enumerate the flux distributions and conduct a differential flux analysis on it.

**Figure 3.**
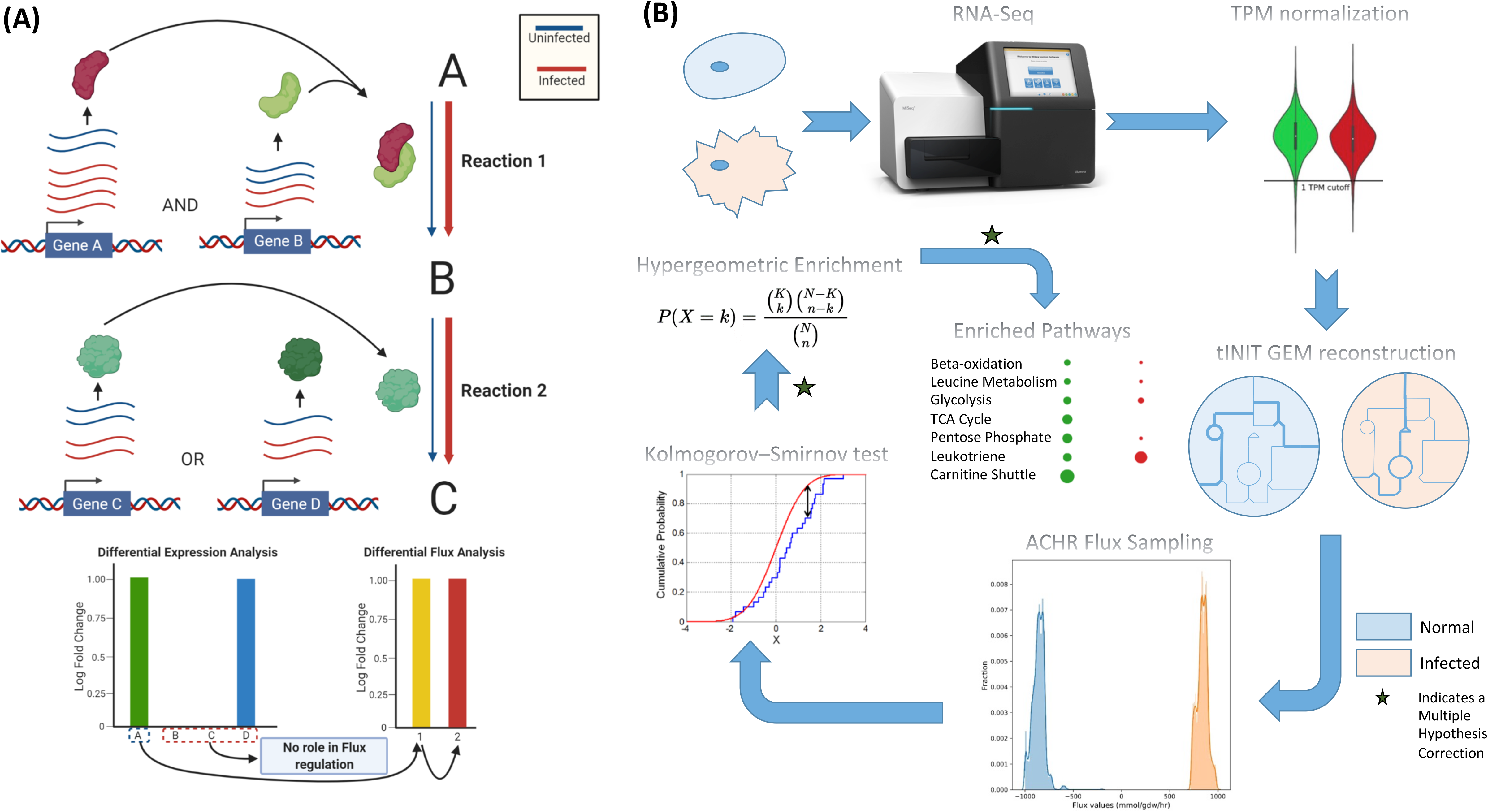
Genome-Scale Differential Flux Analysis (GS-DFA) for finding the metabolic alterations at the level of flux. (A) Boolean relation between activity of enzymes and the gene encoding the proteins in the complex makes the prediction of flux from expression difficult. Gene A and Gene B encodes subunits of the enzyme which catalyzes ‘Reaction 1’. Gene C and Gene D encode two alternative isozymes which catalyzes ‘Reaction 2’. The width of the arrow for ‘Reaction 1’ and ‘Reactions 2’ represents the level of flux. The black arrow in the bottom panel of (A) moves from driver to affected reaction. (B) The pipeline for GS-DFA beginning from RNA-seq of infected and normal cell line, followed by construction of context specific models, flux sampling, reaction filtering and over-representation analysis of pathways. The ‘star’ symbol indicates a multiple hypothesis correction step through Benjamini-Hochberg False Discovery Rate. A two-sample Kolmogorov-Smirnov test is used to differentiate between the probability distribution of flux between diseased and non-diseased state.

Metabolic flux analysis allows us to understand the flux distribution in a model under a given set of condition. While tools like FBA enable us to obtain possible flux values for reactions throughout the metabolic model, it doesn’t guarantee uniqueness of the solution and necessitates an objective function(21). Flux variability analysis and flux sampling allow us to capture the extent of the flux solution space(28). Hence, the information of the solution space can be used to understand the quantitative features of metabolic phenotypes. We were further interested in comparing the flux solution spaces between two conditions (normal vs. diseased) and highlight reactions that have altered flux. But given the lack of precision in metabolic flux analysis, stringent statistical filtering is important for such an analysis.

Flux sampling allows us to screen through all the possible flux values and generate a probability distribution of the values that a given reaction can undertake(29). This can be directed for comparing the flux values distribution between a normal states vs. diseased states. We developed a pipeline (Figure 3B) that takes the context specific models of NHBE and Lung Biopsy samples in both normal and diseased state, performs a differential flux analysis to reveal altered reactions. The pipeline leverages Kolmogorov-Smirnov test to compare the flux probability distribution for given reactions between two conditions. The altered reactions having statistical significant differences in the distributions and a high fold change (F.C. cutoff ~ Flux 1 is 10 times Flux 2) are used for reaction enrichment analysis among the metabolic subsystems present in the model.

For NHBE cell context specific models i.e. iNHBE and iNHBECov2, beta-oxidation of di-unsaturated fatty acids (mitochondria), fatty acid oxidation, fatty acid activation, carnitine shuttle pathway (mitochondria and cytosol), arachidonic acid metabolism, fatty acid elongation and few amino acid biosynthesis pathway were severely altered between the normal vs. diseased state (Figure 4A and 4B). Majority of the altered reactions belonged to fatty acid oxidation pathway. Arachidonic acid metabolism, which has been earlier reported to be one of the severely altered reactions in HCov-229E, also showed altered flux for several reactions(38). Beta-oxidation of fatty acid has also been discussed to play a significant role in the progress of the infection(41). There were several differences in beta-oxidation pathways of di-unsaturated and even-chain fatty acid. Of the amino acid biosynthesis pathways, aromatic amino acid pathway (Phenylalanine, Tyrosine and Tryptophan) was altered at the level of flux between normal and diseased state. Interestingly tryptophan metabolism has been reported to be altered during SARS Cov2 infection through metabolomics studies(15). Alterations in tryptophan metabolism has been implicated in several viral diseases in previous studies and has been proposed as an important therapeutic target. Kynurenine, which is a product derived from tryptophan by the activity of IDO1 (indoleamine 2,3 dioxygenase 1), is an important immune-regulatory molecule. Defects in IDO1 has been shown to be correlated to IL6 induced inflammation in cells(15). There is also significant alterations in nucleotide metabolism. This could results from high demand for RNA synthesis in the host to support the complicated transcriptional readout in the virus and the synthesis of the ssRNA while suppressing the DNA synthesis. Disturbance of arachidonic acid metabolism is very clearly observed through differential flux analysis. An increased activity of carnitine shuttle responsible for fatty acid beta-oxidation is also observed in the altered reactions.

**Figure 4.**
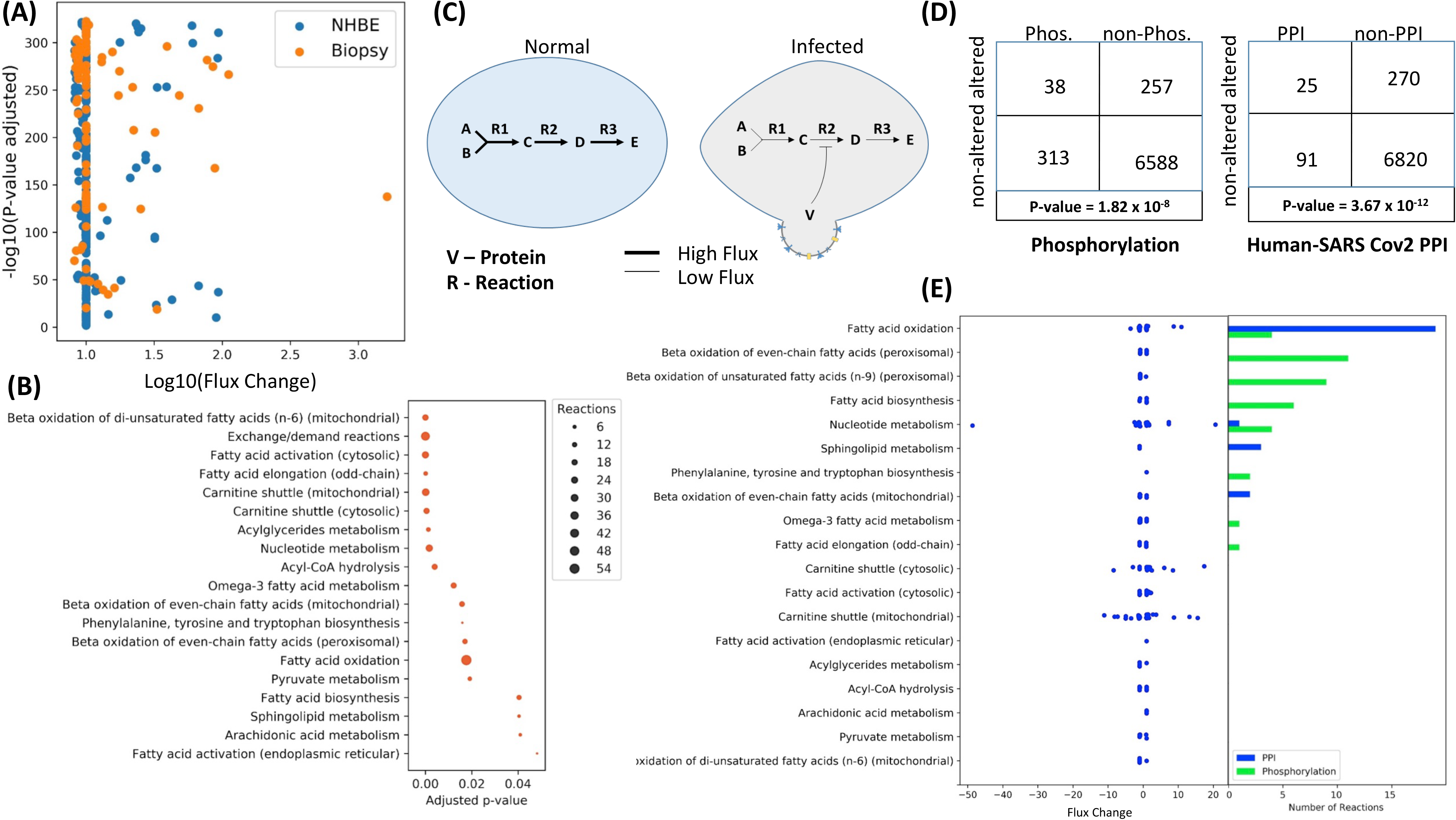
GS-DFA reveals altered pathways in infected cells and post-translational modifications in infected cells are enriched in altered reactions. (A) The volcano-plot for log_10_(flux change) vs ‒log_10_(p-value) for altered reactions. The flux change is calculated as described in materials and methods. (B) Enriched pathways (subsystems) in the SARS Cov2 infected NHBE cells as revealed by GS-DFA analysis. The presented reactions have flux change (as described in materials and methods) greater than 0.82 and have adjusted p-value less than 0.05 (C) The prospective mechanism of flux regulation by post-translational modification or allosteric effects on enzymes mediated by viral proteins. (D) The enrichment of differentially phosphorylated enzymes and enzymes interacting with viral proteins in the set of altered reactions identified by GS-DFA. (E) A pathway wise description of (D) with the flux changes of the constituent altered reactions in the pathways along with the number of such reactions affected by differential phosphorylation and protein-protein interactions with viral proteins.

Several pathways highlighted in this analysis are also the ones which are heavily implicated for progression of diseases of RNA virus if not exactly SARS Cov2. They could serve as potential therapeutic targets. While metabolic flux analysis highlights reactions that are altered based on their flux value distribution, it doesn’t give us any information about the molecular mechanistic principles driving such changes. Thus, it becomes imperative to leverage other ‘layers’ of omics data to understand the molecular modifications on enzymes leading to reduced or enhanced capacity for flux.

### Protein-Protein interactions and altered phosphorylation potentially mediate metabolic changes

It has been reported earlier that most RNA viruses modulate the metabolism and physiology of the host cell through post-translational modifications rather than transcriptional changes(42). We were therefore interested to check if the enzymatic complex driving the reactions affected at the level of flux in diseased state are post-translationally modified. We leveraged the data on binary protein interaction network between host and SARS Cov2 proteins(43), and differential phosphorylation(44) dataset for this purpose. In our proposed model, we suspect several viral proteins interact with metabolic machinery of the host cell and press changes either through i) Allosteric modifications ii) Phosphorylation/Dephosphorylation induced effects (Figure 4C). We did a cross interaction analysis between the altered metabolic pathways and the post-translational modifications (Figure 4D and 4E). Surprisingly, several proteins interaction networks existed between the viral proteins and the enzymes involved in fatty acid oxidation, beta-oxidation of even chain fatty acids, nucleotide metabolism and sphingolipid metabolism. These are some of the severely affected pathways highlighted through our differential flux analysis. Furthermore, overlaying the differential phosphorylation dataset on our metabolic flux analysis, revealed several reactions belonging to beta-oxidation of fatty acids (peroxisomal), fatty acid elongation and omega-3 fatty acid metabolism have modified phosphorylation state between the normal and diseased condition (Figure 4E).

This directly indicates the modification of host metabolism induced by the viral proteins through post-translational modification. We also acknowledge the presence of complicated pathways of modifications through host signaling pathways which we might have missed in our analysis. Phosphorylation and allosteric modification are two significant pathways through which metabolic machinery is regulated at the post-translational level. Inhibitors against the viral proteins mediating such molecular modifications can be used as a therapeutic option to restore metabolic homeostasis in the cell. Nevertheless, fatty acid metabolism appears to be one of the most affected pathways in the host which can be targeted for therapy.

## Discussion

In this study, we propose a computational method for conducting a differential flux analysis of metabolism at a genome-scale. This method allowed us to identify altered reactions in the metabolic network of NHBE cells infected with SARS Cov2 virus. This method comprises of a stringent statistical framework which enables us to filter out reactions which occur by chance due to the fundamental behaviour of linear optimization problems. This method is independent of objective function used and hence can be leveraged for understanding the metabolic flux alterations under given set of conditions in the case of mammalian cells.

Here, we leveraged flux sampling method ACHR (Artificial Coordinate Hit and Run)(29) to sample the flux solution space constrained by thermodynamic bounds and structure of the metabolic network. It is important to understand alternative methods like Flux Variability Analysis(28), which also helps us to probe into the differences in flux distribution in two conditions, assume a uniform distribution of fluxes across the solution space. This assumption makes it useful in allowing us to infer the maximum and minimum flux that a given reaction network can undertake in a given condition. Nevertheless, it fails to consider the likelihood of the actual flux distribution. This adds up to the limitations in inferring precise flux values from genome-scale metabolic reconstructions. Flux sampling analysis by the virtue of random sampling under metabolic and thermodynamic constraints(29, 45), furnishes a probability distribution function for the fluxes in their respective solution space. Thus, we can use statistical tests to obtain confidence intervals of the fluxes and compare them across multiple conditions. This would not have been otherwise possible if we assumed a uniform distribution of flux values in the solution space.

Kolmogorov-Smirnov test compares between the probability distribution of fluxes across several conditions and highlight metabolic modules that are altered. Furthermore, we use multiple-hypothesis testing to ensure the reported significances don’t contain false positives. The enriched set of reactions are further subjected to hypergeometric enrichment test for metabolic subsystems in the model. This is to make sure the reported pathways that are affected are not due to chance. Using this method, we find that several metabolic subsystems affected between diseased and normal state belong to fatty acid pathway in the case of SARS Cov2. Interestingly, there are already several reports which point at the severe deregulation in fatty acid metabolism confirmed at the level of clinical surveys under controlled trails and metabolomics studies(15, 38, 46). In our analysis, we inform pathways such a fatty acid oxidation, arachidonic acid metabolism and beta-oxidation cycles to be severely affected apart from deregulations in amino acid and nucleotide metabolism pathways. The role of beta-oxidation in promoting viral pathogenesis has already been reported in several studies and we believe this could be important for therapeutic design(41). Similarly, disturbances in arachidonic acid metabolism through the linoleic acid-arachidonic acid metabolism axis has been reported to be a hallmark of coronavirus infection(38).

Fatty acid are not only important energy precursors which might be expended by the virus to grow rapidly but also the biochemical precursor for several important signalling molecules. Leukotriene biosynthesis which begins from the oxidation of arachidonic acid has also been inferred to be one of the affected pathways between diseased and normal state. Leukotriene is an important autocrine and paracrine signalling molecule which allows cells to attract immune cells when abnormalities arise is homeostatic state of the cells(30). It is possible that SARS Cov2 strategically interferes with such a signalling pathway by deregulating the biosynthesis pathway for the constituent signalling molecules. We were further interested in understanding how these metabolic changes are imparted on the normal cell by the virus. Metabolic fluxes are always under stringent regulation by the post-translational modifications or allosteric changes in the driver enzyme in shorter time-scales and by transcriptional regulation in longer time-scales. When SARS Cov2 infects the host cell, it is possible that viral proteins interfere with such regulatory controls and this allows the virus to reprogram the metabolism of the cells. A protein-protein interaction network between SARS Cov2 virus and the host cells has been published recently. We leveraged the same to check if the interactions of viral proteins with host proteins are enriched in the set of reactions we inferred from differential flux analysis. Surprisingly, we found several protein interactions and also phosphorylation changes in the enzymes subunits that are associated with altered flux. To infer the impact of protein interactions and phosphorylation on the enzyme functions, we used the Boolean relation between the enzyme subunits and the resulting functionality.

We consider this method to be a significant advancement in the field of metabolic flux analysis. This would allow us to probe into differences in flux apart from differences in network structure among several conditions. This would help us to design experiments to measure the effects of infections on specific metabolic modules in the cell. Such an analysis along with other multi-omics datasets like metabolomics, phospho-proteomics and interactomics will allow us to understand the molecular mechanistic principles of infections. Integration of multi-omics data is expected to reveal several mechanistic phenomena of diseases and will help us design better therapeutics leveraging the power of systems biology.

## Materials and Methods

The workflow for the construction of SARS Cov2 viral biomass equation is mentioned in SI Appendix, SI Materials and Methods. The procedure for building the context specific models of normal and SARS Cov2 infected NHBE and Human Lung cells is described in SI Appendix, SI Materials and Methods. The pipeline for conducting the GS-DFA is also described in SI Appendix, SI Materials and Methods. The integrated analysis of phosphor-proteomics and virus-host protein-protein interactions is mentioned in SI Appendix, SI Materials and Methods.

## Data availability

All the data generated in this study has been provided in SI appendix, Dataset S1-S2. All the codes used for the analysis is available on GitHub (https://github.com/piyushnanda-sysbio/GS-DFA.git).

## Acknowledgments

Authors are thankful to Department of Science and Technology (Grant No. ECR/2016/001096), Department of Biotechnology (Grant No. BT/RLF/Re-entry/06/2013) and Scheme for Promotion of Academic and Research Collaboration (SPARC), MHRD, Govt. of India (Grant No. SPARC/2018-2019/P265/SL).

## Notes

### Competing Interest Statement

The authors have declared no competing interest.

## References

1. B. H. Goodpaster, L. M. Sparks, Metabolic Flexibility in Health and Disease. Cell Metab. 25, 1027–1036 (2017).

2. E. Holmes, I. D. Wilson, J. K. Nicholson, Metabolic Phenotyping in Health and Disease. Cell 134, 714–717 (2008).

3. P. Kandasamy, G. Gyimesi, Y. Kanai, M. A. Hediger, Amino acid transporters revisited: New views in health and disease. Trends Biochem. Sci. 43, 752–789 (2018).

4. E. Tzika, T. Dreker, A. Imhof, Epigenitics and metabolism in health and disease. Front. Genet. 9 (2018).

5. L. Yang, J. T. Yurkovich, Z. A. King, B. O. Palsson, Modeling the multi-scale mechanisms of macromolecular resource allocation. Curr. Opin. Microbiol. 45, 8–15 (2018).

6. C. Zhang, Q. Hua, Applications of genome-scale metabolic models in biotechnology and systems medicine. Front. Physiol. 6 (2016).

7. S. L. Berger, P. Sassone-Corsi, Metabolic signaling to chromatin. Cold Spring Harb. Perspect. Biol. 8, 1–23 (2016).

8. S. Trefely, C. D. Lovell, N. W. Snyder, K. E. Wellen, Compartmentalised acyl-CoA metabolism and roles in chromatin regulation. Mol. Metab. 38, 100941 (2020).

9. S. Sivanand, I. Viney, K. E. Wellen, Spatiotemporal Control of Acetyl-CoA Metabolism in Chromatin Regulation. Trends Biochem. Sci. 43, 61–74 (2018).

10. I. Martínez-Reyes, N. S. Chandel, Mitochondrial TCA cycle metabolites control physiology and disease. Nat. Commun. 11, 1–11 (2020).

11. M. Neagu, et al., Inflammation and metabolism in cancer cell—mitochondria key player. Front. Oncol. 9 (2019).

12. F. Hirschhaeuser, U. G. A. Sattler, W. Mueller-Klieser, Lactate: A metabolic key player in cancer. Cancer Res. 71, 6921–6925 (2011).

13. B. M. Cumming, K. W. Addicott, J. H. Adamson, A. J. C. Steyn, Mycobacterium tuberculosis induces decelerated bioenergetic metabolism in human macrophages. Elife 7 (2018).

14. V. R. Muddapu, S. A. P. Dharshini, V. S. Chakravarthy, M. M. Gromiha, Neurodegenerative Diseases – Is Metabolic Deficiency the Root Cause? Front. Neurosci. 14 (2020).

15. T. Thomas, et al., COVID-19 infection alters kynurenine and fatty acid metabolism, correlating with IL-6 levels and renal status. JCI Insight 5 (2020).

16. A. Renz, L. Widerspick, A. Dräger, D. Dräger, FBA reveals guanylate kinase as a potential target for antiviral therapies against SARS-CoV-2. 1–11 (2020).

17. M. Abu-Farha, et al., The role of lipid metabolism in COVID-19 virus infection and as a drug target. Int. J. Mol. Sci. 21 (2020).

18. S. Chandrasekaran, N. D. Price, Probabilistic integrative modeling of genome-scale metabolic and regulatory networks in Escherichia coli and Mycobacterium tuberculosis. Proc. Natl. Acad. Sci. U. S. A. 107, 17845–17850 (2010).

19. N. C. Duarte, et al., Global reconstruction of the human metabolic network based on genomic and bibliomic data. Proc. Natl. Acad. Sci. U. S. A. 104, 1777–1782 (2007).

20. O. Folger, et al., Predicting selective drug targets in cancer through metabolic networks. Mol. Syst. Biol. 7 (2011).

21. S. Opdam, et al., A Systematic Evaluation of Methods for Tailoring Genome-Scale Metabolic Models. Cell Syst. 4, 318–329.e6 (2017).

22. E. Brunk, et al., Recon3D enables a three-dimensional view of gene variation in human metabolism. Nat. Biotechnol. 36, 272–281 (2018).

23. J. L. Robinson, et al., An atlas of human metabolism. Sci. Signal. 13, 1–12 (2020).

24. J. D. Orth, I. Thiele, B. O. Palsson, What is flux balance analysis? Nat. Biotechnol. 28, 245–248 (2010).

25. V. Pandey, N. Hadadi, V. Hatzimanikatis, Enhanced flux prediction by integrating relative expression and relative metabolite abundance into thermodynamically consistent metabolic models. PLoS Comput. Biol. 15 (2019).

26. K. Yizhak, T. Benyamini, W. Liebermeister, E. Ruppin, T. Shlomi, Integrating quantitative proteomics and metabolomics with a genome-scale metabolic network model. Bioinformatics 26 (2010).

27. A. Mardinoglu, F. Gatto, J. Nielsen, Genome-scale modeling of human metabolism - a systems biology approach. Biotechnol. J. 8, 985–996 (2013).

28. R. Mahadevan, C. H. Schilling, The effects of alternate optimal solutions in constraint-based genome-scale metabolic models. Metab. Eng. 5, 264–276 (2003).

29. S. Bordel, R. Agren, J. Nielsen, Sampling the solution space in genome-scale metabolic networks reveals transcriptional regulation in key enzymes. PLoS Comput. Biol. 6, 16 (2010).

30. C. D. Funk, Prostaglandins and leukotrienes: Advances in eicosanoid biology. Science (80-.). 294, 1871–1875 (2001).

31. Y. M. Bar-on, A. Flamholz, R. Phillips, R. Milo, SARS-CoV-2 (COVID-19) by the numbers. Elife (2020).

32. N. Irigoyen, et al., High-Resolution Analysis of Coronavirus Gene Expression by RNA Sequencing and Ribosome Profiling. PLoS Pathog. 12 (2016).

33. X. Xia, Extreme Genomic CpG Deficiency in SARS-CoV-2 and Evasion of Host Antiviral Defense. Mol. Biol. Evol. 37, 2699–2705 (2020).

34. A. C. Walls, et al., Glycan shield and epitope masking of a coronavirus spike protein observed by cryo-electron microscopy. Nat. Struct. Mol. Biol. 23, 899–905 (2016).

35. N. S. Ogando, et al., SARS-coronavirus-2 replication in Vero E6 cells: Replication kinetics, rapid adaptation and cytopathology. J. Gen. Virol. 101, 925–940 (2020).

36. R. Agren, et al., Reconstruction of genome-scale active metabolic networks for 69 human cell types and 16 cancer types using INIT. PLoS Comput. Biol. 8 (2012).

37. D. Blanco-Melo, et al., Imbalanced Host Response to SARS-CoV-2 Drives Development of COVID-19. Cell 181, 1036–1045.e9 (2020).

38. B. Yan, et al., Characterization of the lipidomic profile of human coronavirus-infected cells: Implications for lipid metabolism remodeling upon coronavirus replication. Viruses 11 (2019).

39. D. Bojkova, et al., Proteomics of SARS-CoV-2-infected host cells reveals therapy targets. Nature 583, 469–472 (2020).

40. J. S. Kang, Dietary restriction of amino acids for Cancer therapy. Nutr. Metab. 17 (2020).

41. J. S. Ayres, A metabolic handbook for the COVID-19 pandemic. Nat. Metab. 2, 572–585 (2020).

42. G. A. Gualdoni, et al., Rhinovirus induces an anabolic reprogramming in host cell metabolism essential for viral replication. Proc. Natl. Acad. Sci. U. S. A. 115, E7158–E7165 (2018).

43. D. E. Gordon, et al., A SARS-CoV-2 protein interaction map reveals targets for drug repurposing. Nature 583, 459–468 (2020).

44. M. Bouhaddou, et al., The Global Phosphorylation Landscape of SARS-CoV-2 Infection. Cell 182, 685–712.e19 (2020).

45. J. Schellenberger, B. Palsson, Use of randomized sampling for analysis of metabolic networks. J. Biol. Chem. 284, 5457–5461 (2009).

46. Q. Wu, et al., Altered Lipid Metabolism in Recovered SARS Patients Twelve Years after Infection. Sci. Rep. 7 (2017).

