## Supplementary Information for "Genome Scale-Differential Flux Analysis reveals deregulation of lung cell metabolism on SARS Cov2 infection"

### Supplementary Information Text

### Materials and Methods:

#### Stoichiometric reconstruction of Virus Biomass Objective Function (VBOF) for SARS Cov2

All the calculations for nucleic acids, proteins, carbohydrates and lipids are conducted per gram dry weight (GDW) of the virus. The nucleotide content was calculated from the single stranded positive RNA sequence which is present as a single copy. The contribution of transcriptome towards nucleotide was also considered. Briefly, the number of transcripts per gram dry weight of the virus was calculated from the proteomics and translational efficiency data(1). The translational efficiency is defined as the number of mRNA transcripts which are converted to polypeptides. The translational efficiency of various proteins were in the range ~ 0.1 to 2(1). The highest efficiency was for the N protein while the lowest was for ORF1A and ORF3A. Nucleotides were the principal components of the biomass equation due to the high demands for mRNA synthesis apart from viral genome synthesis.

The contribution of amino acids to viral biomass was calculated from a proteomics dataset(2) published earlier. The copy number of spike (S) protein was considered as ~300 subunits per virion and the absolute stoichiometry of other proteins were calculated using it as a basis. It was important to validate the copy number of S protein on SARS Cov2. While it has been reported to approximately be the same as that for SARS Cov(3), it was imperative for us to validate it as VBOF calculation is directly dependent on that. We therefore leveraged electron micrographs of SARS Cov2 and used image analysis to estimate the copy number of spike protein on each virion. (Figure S1 and Extended Methods).

N-acetyl Glucosamine (NAG) content of the biomass was calculated from the PDB file (PDB ID: 5SZS) of S protein. Briefly, the number of NAG molecules present in the crystal structure of a single S protein subunit was calculated. We obtained 63 NAG molecules per subunit of the virus in this way. The total number of NAG molecules studded on the S protein was calculated using the copy number information obtained above. The lipid composition was derived from previous studies(4). Briefly, the total molecules of lipids present on the viral membrane was calculated from the surface area assuming 2 molecules per $nm^2$ as discussed in earlier studies(5).

The total virus molecular weight was calculated from the sum of mass of all macromolecules present in the virus (Dataset S1). The units of all the biomass precursors were converted to millimoles per GDW of SARS Cov2 using this information. The reconstructed biomass equation i.e. Cov2VBOF (SARS Cov2 Virus Biomass Objective Function) was added to HumanGEM model. Appropriate exchange reactions were added to enable flux through the VBOF.

### Integration of gene expression data using tINIT

Gene expression data normalization

The gene expression data was derived from GSE147507. The triplicate data for mock infected NHBE cells and SARS Cov2 infected NHBE cells was considered. The information regarding the transcript isoform was obtained from Mammalian Transcript Database (For Bronchial Epithelial Cells). The dataset was normalized to TPM (Transcripts per Million reads) using the transcript lengths specific for NHBE cells (Dataset S2). Custom scripts were written in R for the same.

tINIT for integration of gene expression

The latest version of tINIT (version 2.0) was used to integrate the gene expression data into HumanGEM model(6). The steps for reconstruction has been described in Figure S2. Briefly, the highly curated HumanGEM model in a closed form i.e. all exchange reactions closed was fed to the program "gettINITModel2" in MATLAB 2017b with COBRA Toolbox v3.0(7) along with the TPM normalized gene expression data. A gene expression cutoff of 1 TPM was used for the algorithm as reported earlier(6). The integer optimization algorithm underlying tINIT tries to minimize the incorporation of transcripts below this level. The task list for tINIT was prepared as reported earlier(6). The task list for the creation of context specific model of NHBE cells infected with SARS Cov2 has additional reactions (Dataset S2). Apart from synthesizing non-essential amino acids, conduct oxidative phosphorylation and other basic metabolic tasks, the model was expected to conduct specialized tasks necessary for the growth of the virus. These additional reactions were the tasks to de-novo synthesize NAG needed for the synthesis of viral biomass and produce viral biomass i.e. Cov2VBOF from the components in the media (Dataset S2). IBM CPLEX 12.1 (IBM Academic License) was used for running the tINIT program.

The resulting context specific models was constrained using the uptake rates from media components as reported in several literature (Dataset S2). Flux Balance Analysis using either the HumanGEM biomass reaction or Cov2VBOF was used to validate the doubling time of the NHBE cells and SARS Cov2 respectively. This was compared to the experimentally measured data and a t-test was conducted to show no significant differences between the experimental and theoretical growth rate for SARS Cov2.

Henceforth, we will call the context specific model for normal NHBE cells and SARS Cov2 infected NHBE cells as iNHBE and iNHBECov2.

VBOF sensitivity analysis

It was important check for the sensitivity of the specific growth rate prediction with respect to variations in coefficient (mmoles/gdw) or fractional composition of biomass precursors. We varied the coefficients of biomass precursors one at a time by ±10% and calculated the growth rate by flux balance analysis. The FBA was set-up on the modified SARS Cov2 biomass equations with coefficients increased by ±10% one at a time as follows:

$$\begin{aligned} &maximize\ c*^T v \\ &subject\ to\ i)\ S.v = 0 \\ &\qquad ii)\ lowerbound \le v \le upperbound \end{aligned}$$

Here, c* is the modified biomass coefficient vector such that **$c^* = (c_1, c_2, c_3 \ldots c_n \pm 0.1c_n)$.** Here, the coefficient for $c_n$ biomass precursor has been manipulated by ±10% whereas the coefficient of other biomass precursors remain the same.

IBM CPLEX solver was used to conduct the flux balance analysis. The context-specific model of NHBE infected with SARS Cov2 i.e. iNHBECov2 was considered for the analysis constrained with HAM media.

**Differential Flux Analysis**

Flux sampling

ACHR (Artificial Co-ordinate Hit and Run) was used to sample fluxes from iNHBE and iNHBECov2. A thinning factor of 100 was used to obtain sparse and un-correlated fluxes which helps to span the whole flux space. IBM CPLEX was used for the optimization underlying ACHR flux sampling. 10,000 sample points were drawn for each model from the sampling object. Custom scripts were written in Python 3.7 for the flux sampling estimation.

Determining differentially altered flux

Two-sample Kolmogorov Smirnov Test (KS Test) was used to differentiate between the flux distribution between iNHBE and iNHBECov2 models. A significance level of 0.05 was used for filtering the altered reactions. Moreover, we calculated the change in flux between two models from the sample means of the distributions. The flux mean in the case of infected and uninfected cells is the statistical mean of the flux distribution of each. $S_{infected}$ and $S_{uninfected}$ are the flux distributions for a given reactions in infected and uninfected conditions. The bar indicates the arithmetic mean of the distribution which is the most represented flux from the distributions. The flux change was calculated for each reaction as follows:

$$Flux\ Change\ (FC) = \frac{\bar{S}_{infected} - \bar{S}_{uninfected}}{|\bar{S}_{infected} + \bar{S}_{infected}|}$$

The denominator is used to normalize the magnitude of flux change. This is avoid biased filtering out of reactions with low flux but of important biological consequence. A Flux

Change (FC) of 0.82 (corresponding to 10 fold change in flux) was used to filter reactions with insignificant flux change. A Benjamini-Hochberg FDR Multiple hypothesis correction with α=0.05 was used to correct the p-values for all the resulting differentially altered reactions. The adjusted p-value was used along with the FC cutoff of 0.82 for filtering out the reactions which would also occur by chance.

Reaction Enrichment Analysis

Hypergeometric enrichment was used to obtain HumanGEM subsystems(8) (Dataset S2) which are overrepresented in the altered set of reactions. Briefly, the enriched reaction list obtained from above was used to conduct a two-tailed hypergeometric test. Here, adjusted p-value was used to obtain the pathways showing significant representation based on the constituent altered reactions.

$$P(X = k) = \frac{\binom{K}{k}\binom{N-K}{n-k}}{\binom{N}{n}}$$

Here, P(X=k) is the probability that we find k reactions just by chance in a given subsystem. K is the total number of reactions belonging to a given subsystem, N is the total number of reactions in the model and n is the total number of reactions that show altered flux as calculated by differential flux analysis. For each subsystem in the model, we conducted a hypergeometric enrichment test for overrepresentation. If P(X=k)<0.05, then it is likely that the over-representation of the subsystem is due to the high number of altered reactions in the pathway rather than by chance. The resulting p-values for pathways were again subjected to multiple-hypothesis correction using Benjamini-Hochberg method using False Discovery Rate (FDR) with α=0.05. From this, we were able to obtain the pathways in the HumanGEM model which are most affected by the infection.

**Integrated Analysis of Protein-Protein Interaction Network (PPINs) and Differential Phosphorylation (DPs) with altered reactions**

Reactions affected by PPINs and DPs

The PPIN dataset was obtained from recently published literature on host-virus interactome(9). It is to be noted that the PPIN screen was conducted in a different cell line i.e. HEK-293T/17 cells. The differential phosphorylation data was obtained from another recently published report on phosphoproteomics(10). Here, the datasets were either the interaction of viral proteins or phosphoproteome with the human gene products not with the protein complex. The association of the gene product with the activity of the protein complex can be presented by Boolean relation(11). Similarly, in the case of metabolic model, the relationships between the gene product and the activity of enzyme is represented by a Boolean logic i.e. GPR (Gene-Protein-Reaction) association. Briefly, an OR relationship between two gene products represents isozymes i.e. Two alternative gene product that can furnish the same activity. An AND relationship between two gene products implies that both the gene products are required to form an active enzyme complex. In order to map the effect of PPINs and DPs on the reactions, it was necessary to consider the Boolean relation. We used mapExpressionToReaction () within COBRA Toolbox v3.0 in MATLAB 2017b to map the effect on the gene product to the effect on the reactions. The gene products which were either associated with PPINs or DPs, we assign a value of 0 and the other non-interacting gene products were assigned a value of 1. If an enzyme is formed by complementation of two gene products i.e. AND relation, the reaction is affected even when just one of the gene products is associated with PPINs or DPs. But if the enzymatic activity is alternatively fulfilled by two separate gene products, the reaction is affected only when both of the gene products are associated with PPINs or DPs. Here, we consider PPINs and DPs are processes that influence activity of the gene products either through allosteric modifications or post-translational modifications. It is a characteristic nature of RNA virus to manipulate metabolism and regulation in the host mainly through short-term PPINs or other modifications rather than heavy transcriptional alteration. mapExpressionToReaction () yield a score for each reaction based on whether the reaction is affected (0) or unaffected(1) by the PPINs and DPs.

This set of reactions was compared to the set of reactions which were found to be differential altered in term of flux. A hypergeometric test was conducted to check if the enrichment of PPINs and DPs affected reactions in the set of reactions having differential flux were due to chance or not. The hypergeometric test was conducted as follows:

$$P(X = k) = \frac{\binom{K}{k}\binom{N-K}{n-k}}{\binom{N}{n}}$$

Here, P(X=k) is the probability that we find k reactions affected by PPINs or DPs just by chance in a given subsystem. K is the total number of reactions affected by PPINs or DPs, N is the total number of reactions in the model and n is the total number of reactions that show altered flux as calculated by differential flux analysis. For each subsystem in the model, we conducted a hypergeometric enrichment test for overrepresentation. If $P(X=k) < 0.05$, then it is likely that the over-representation of PPINs or DPs affected reactions are due to real effects of PPIN and DP on metabolism rather than by chance.

**Extended Methods:**

Estimating spike protein stoichiometry from electron microscopy images

The calculation of absolute protein numbers for SARS Cov2 is entirely based on the number of spike protein subunits on the virus. In recent discussions on the chemical composition of the virus(3), the spike protein subunit counts on SARS Cov2 has been assumed to be the same as that of SARS Cov. While this doesn't entirely defy logic, it would be imperative to validate the protein counts per virus in the SARS Cov2. For this purpose, we leveraged the electron micrographs of the virus taken by various research organizations. We used a combination of image analysis and mathematical modeling to derive the number of spike protein per virus.

Briefly, the electron micrographs were converted to 16 bit images. It could be observed that in several electron micrographs, the spike protein appear as the protrusions around the virus. We measured the intensities of multiple-points along the circumference of the virus. The number of spike proteins will be roughly equal to the number of positions along the circumference where we see high intensity values. We used ImageJ (FIJI) to estimate the intensity along the circumference of the virus at roughly uniformly placed points in order. It is to be noted that each intact spike protein comprises of 3 subunits. The plot between the intensity of various points (y-axis) and the angular position (x-axis) would give us intensity distribution at various angular positions. The number of peaks (corresponding to high intensity spots) would be roughly equal to the number of spike proteins (trimers) on the surface. We used custom codes written in MATLAB and leveraged 'findpeaks' functions to get the number of such peaks. This gave us the distribution of spike proteins (spike protein counts per circumference) in 2 dimensional cross section of the virus.

In order to estimate the distribution of spike proteins in the 3D surface of the virus, we assumed a uniform distribution of the spike protein on the surface. The following derivation was used to calculate the spike protein distribution (spike protein count per virus) on the surface:

N= Number of spike proteins calculated from the electron micrograph i.e. spike proteins along the circumference

n= Spike proteins per unit length circumference = $\frac{N}{2 \times \pi \times R}$

R = Radius of the cross section of the virus

Consider an elemental disk of width dx at a distance x from the center of the sphere.

Total number of spike protein on the elemental disk = *dn* = Spike proteins along the circumference of the disk x Spike protein along the width of the disk

$$dn = \left[\frac{N}{2 \times \pi \times R} \times 2\times\pi \times\sqrt{R^2 - x^2}\right] \times \left[\frac{N}{2 \times \pi \times R} \times dx\right]$$

The integration of *dn* while x increases from 0 to R will give us the distribution of spike proteins in one hemisphere of the circle.

Therefore,

Let $N_{total}$ = Total number of spike protein on the surface

$$0.5 \times N_{total} = \int_0^R dn$$

$$0.5 \times N_{total} = \int_0^R \left[\frac{N}{2 \times \pi \times R} \times 2 \times \pi \times \sqrt{R^2 - x^2}\right] \times \left[\frac{N}{2 \times \pi \times R} \times dx\right]$$

$$0.5 \times N_{total} = \frac{N^2}{8}$$

$$N_{total} = \frac{N^2}{4} \quad ..(i)$$

Figure S1 shows the number of spike proteins calculated from the electron micrographs (N). The average N i.e. <N> ~ 20. Substituting that in (i), we get $N_{total}$ ~ 100. Note that $N_{total}$ is the total number of spike trimers, so the total number of spike subunits on the surface is ~300 which is the same reported for SARS Cov in recent discussions.

Hence, this analysis provides direct evidence of spike protein counts on the surface of the virus and validates the assumption. We can therefore use this estimated protein count for generation of biomass objective function/biomass equation for the SARS Cov2 virus.

**Figure captions:**

**Figure S1**: The electron micrograph images of SARS Cov2 and calculation of spike protein count per virion from it. A) The electron micrograph along with the contributing organization. B) Number of spike proteins calculated from intensity measurements of the micrograph along the circumference of the virus.

**Figure S2**: Pipeline for processing of RNA Seq data and tINIT for integration of gene expression data into the model. The processed final model is used for differential flux analysis.

**Figure S3**: Effect of change in uptake rates of nutrient on the fitness of the virus and NHBE cells. The uptake rates were varied and a point optimization program was used to calculate the specific growth rate. The fitness change is reported as the ratio of specific growth rate under perturbation to the specific growth rate under normal uptake rate. All the uptake rates were negative.

**Figure S4**: Sensitivity analysis of biomass objective function (VBOF) with respect to variations in biomass composition. The coefficients of biomass precursors were varied by ±10% taken one at a time and the specific growth rate was calculated by FBA.

**Supplementary datasets:**

**Dataset S1:** The reconstruction of SARS Cov2 biomass equation from the stoichiometric composition of the virus. The sheets in the dataset comprises of stepwise calculation of SARS Cov2 amino acid composition, nucleotide composition, carbohydrate and lipid composition. The last sheet comprises of the final viral biomass equation.

**Dataset S2:** The datasets in S2 relate to various input files required for model reconstruction, the raw results from the analysis and other details. The description of each dataset within Dataset S2 are given in the first sheet.

### References


1. N. Irigoyen, *et al.*, High-Resolution Analysis of Coronavirus Gene Expression by RNA Sequencing and Ribosome Profiling. *PLoS Pathog.* **12** (2016).
2. D. Bojkova, *et al.*, Proteomics of SARS-CoV-2-infected host cells reveals therapy targets. *Nature* **583**, 469–472 (2020).
3. Y. M. Bar-on, A. Flamholz, R. Phillips, R. Milo, SARS-CoV-2 ( COVID-19 ) by the numbers. *Elife* (2020).
4. G. Van Meer, Lipids of the Golgi membrane. *Trends Cell Biol.* **8**, 29–33 (1998).
5. B. Brügger, *et al.*, The HIV lipidome: A raft with an unusual composition. *Proc. Natl. Acad. Sci. U. S. A.* **103**, 2641–2646 (2006).
6. J. L. Robinson, *et al.*, An atlas of human metabolism. *Sci. Signal.* **13** (2020).
7. Laurent Heirendt, *et al.*, Creation and analysis of biochemical constraintbased models using the COBRA Toolbox v.3.0. *Nat. Protoc.* **14**, 639–702 (2018).
8. J. L. Robinson, *et al.*, An atlas of human metabolism. *Sci. Signal.* **13**, 1–12 (2020).
9. D. E. Gordon, *et al.*, A SARS-CoV-2 protein interaction map reveals targets for drug repurposing. *Nature* **583**, 459–468 (2020).
10. M. Bouhaddou, *et al.*, The Global Phosphorylation Landscape of SARS-CoV-2 Infection. *Cell* **182**, 685-712.e19 (2020).
11. S. Opdam, *et al.*, A Systematic Evaluation of Methods for Tailoring Genome-Scale Metabolic Models. *Cell Syst.* **4**, 318-329.e6 (2017).

**Figure S1**

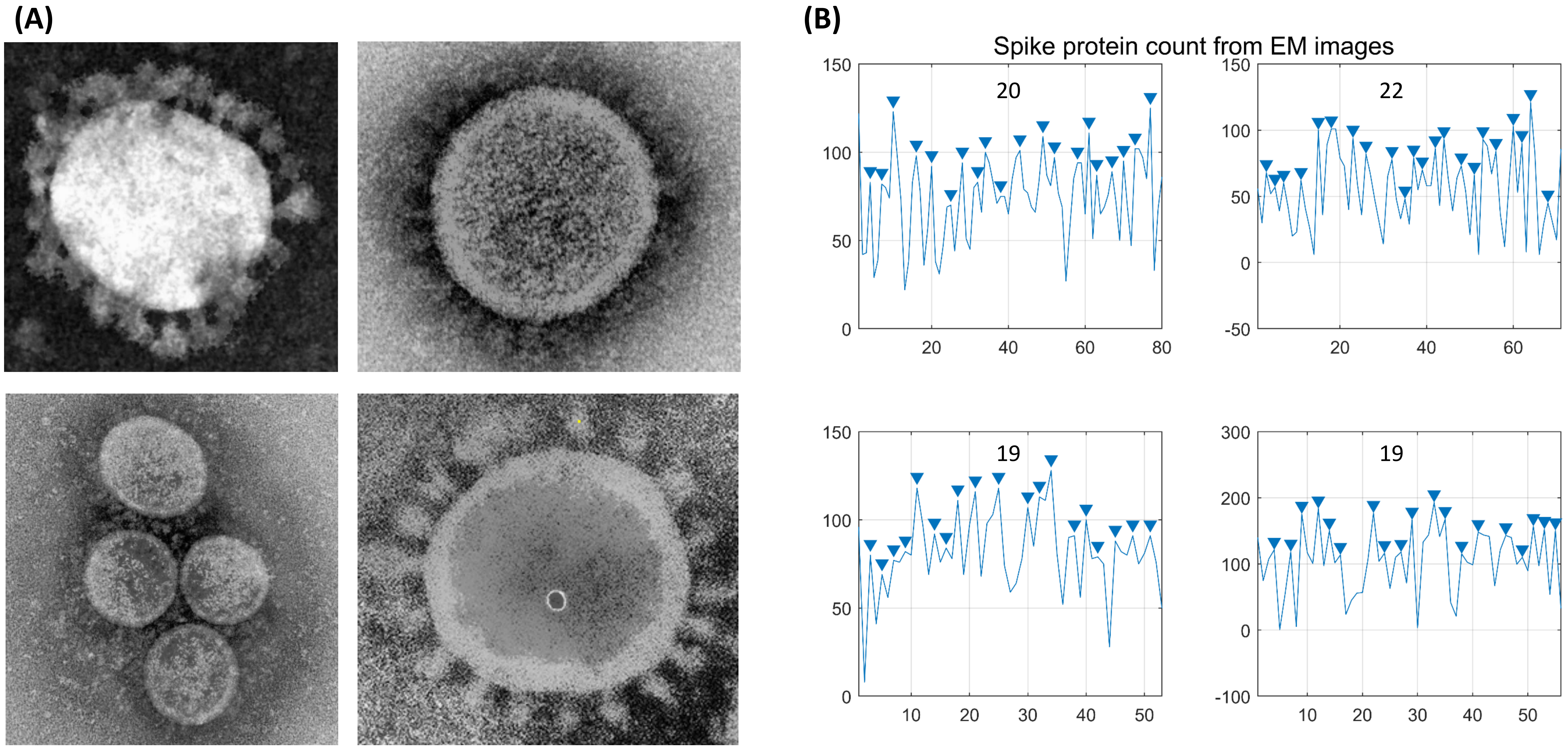
(A)
(B)
Spike protein count from EM images
20
22
19
19
150
100
50
0
-50
-100
200
300
10
20
30
40
50
60
80

**Figure S2**

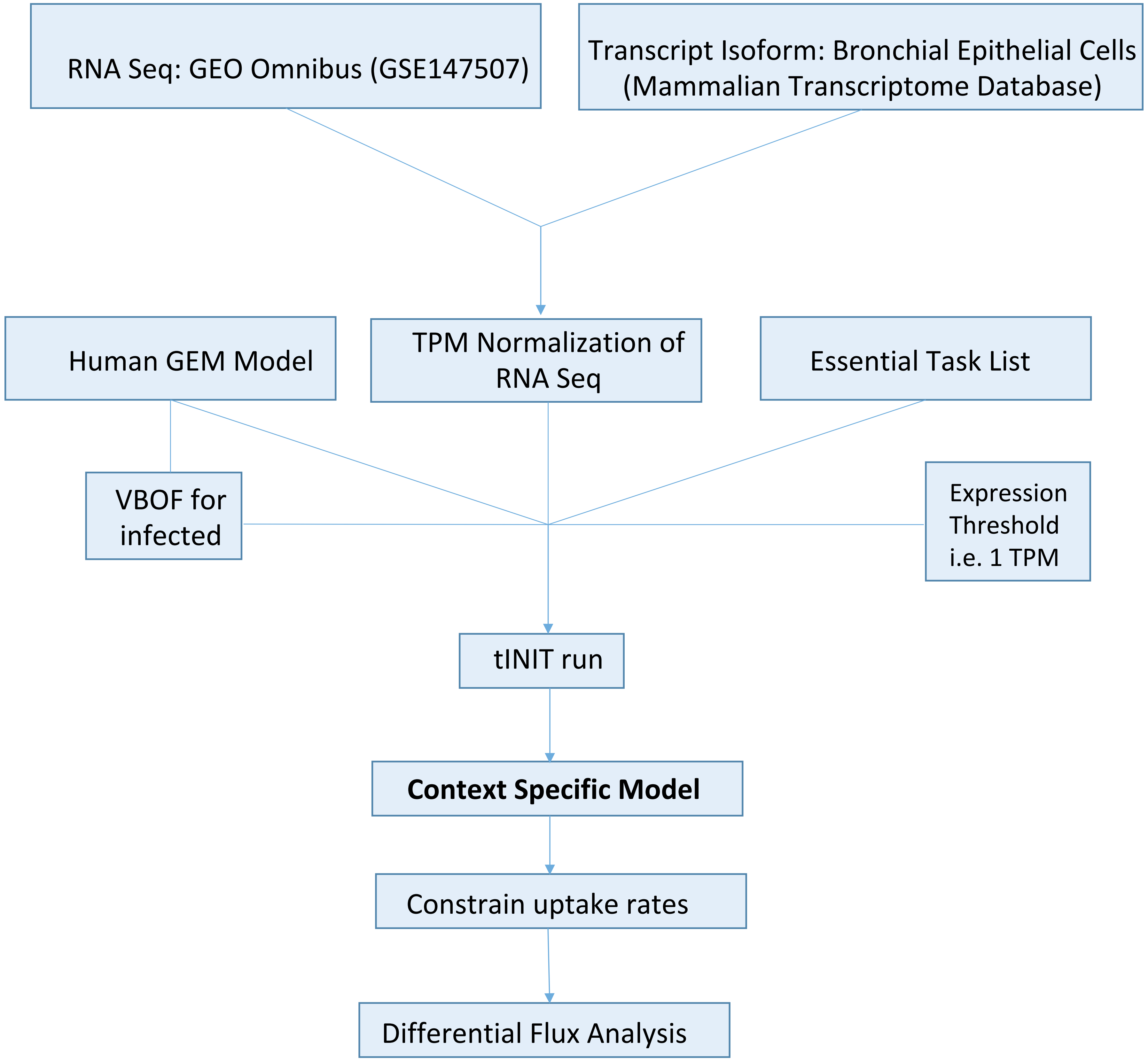
RNA Seq: GEO Omnibus (GSE147507)
Transcript Isoform: Bronchial Epithelial Cells (Mammalian Transcriptome Database)
Human GEM Model
TPM Normalization of RNA Seq
Essential Task List
VBOF for infected
Expression Threshold i.e. 1 TPM
tINIT run
Context Specific Model
Constrain uptake rates
Differential Flux Analysis

### Figure S3

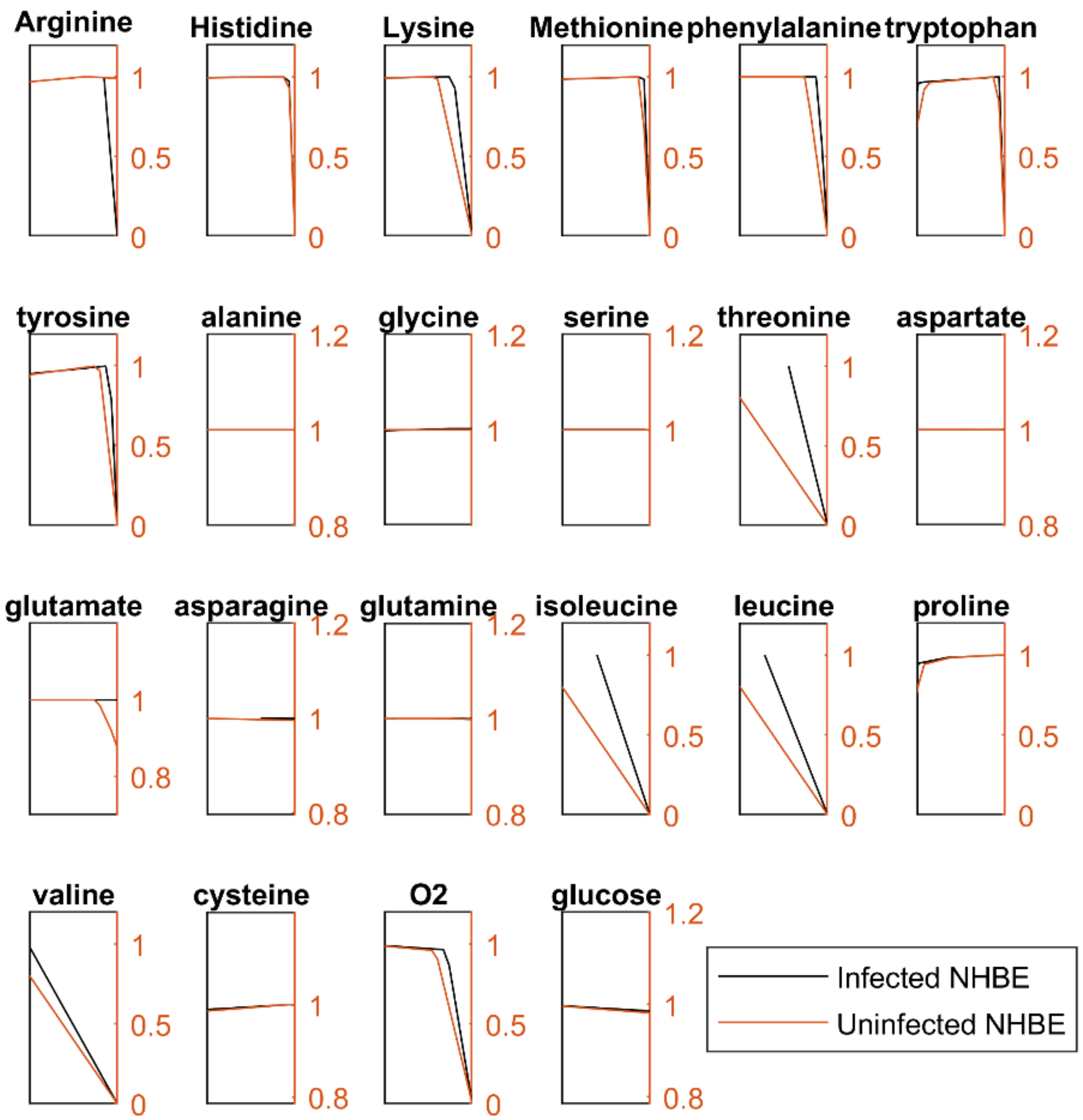
Arginine
Histidine
Lysine
Methionine
phenylalanine
tryptophan
tyrosine
alanine
glycine
serine
threonine
aspartate
glutamate
asparagine
glutamine
isoleucine
leucine
proline
valine
cysteine
O2
glucose
Infected NHBE
Uninfected NHBE

**Figure S4**

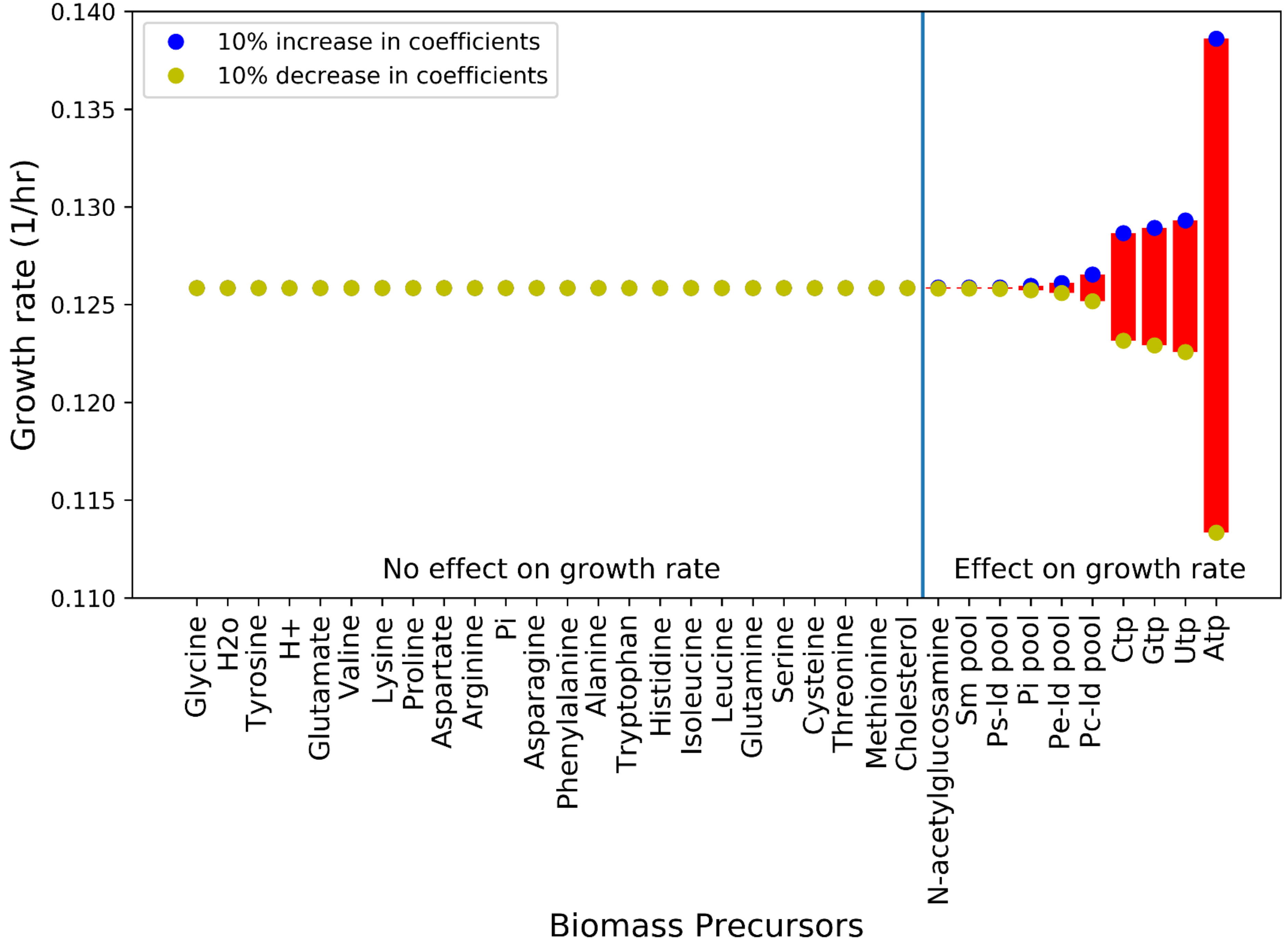
10% increase in coefficients
10% decrease in coefficients
Growth rate (1/hr)
0.140
0.135
0.130
0.125
0.120
0.115
0.110
No effect on growth rate
Effect on growth rate
Glycine
H2o
Tyrosine
H+
Glutamate
Valine
Lysine
Proline
Aspartate
Arginine
Pi
Asparagine
Phenylalanine
Alanine
Tryptophan
Histidine
Isoleucine
Leucine
Glutamine
Serine
Cysteine
Threonine
Methionine
Cholesterol
N-acetylglucosamine
Sm pool
Ps-ld pool
Pi pool
Pe-ld pool
Pc-ld pool
Ctp
Gtp
Utp
Atp
Biomass Precursors
